# Notch as a Driver of Lineage Plasticity and Therapeutic Target in Enzalutamide-Resistant Prostate Cancer

**DOI:** 10.1101/2025.05.26.656166

**Authors:** Yuyin Jiang, Siyuan Cheng, Longjun Li, Melanie Fraidenburg, Isaac Yi Kim, Su Deng, Ping Mu

**Author notes:** Correspondence to (P.M.), (S.D.).

## Abstract

Resistance to androgen receptor (AR)-targeted therapies, such as enzalutamide, in castration-resistant prostate cancer (CRPC) remains a significant clinical challenge, often driven by mechanisms including lineage plasticity. The precise molecular mechanisms driving this process, particularly downstream effectors, remain incompletely understood. Given its established roles in cell fate and stemness, alongside its complex functions in prostate cancer, the Notch signaling pathway presented a compelling focus for study. This study investigates the role of Notch signaling in mediating lineage plasticity and therapeutic resistance in CRPC. Employing transcriptomic analysis and functional assays, we identified Notch activity is elevated across prostate cancer progression resistance. Notably, both CRISPR-mediated knockout and targeted inhibition of Notch reversed enzalutamide resistance *in vitro*. Collectively, this study delineates dynamic alterations in Notch signaling activity during prostate cancer progression and establishes its function as a crucial and druggable driver of therapy resistance. These findings underscore Notch signaling as a promising therapeutic target to counteract resistance to AR-targeted therapies in advanced prostate cancer.

## Introduction

Prostate cancer remains a leading malignancy in men and is fundamentally driven by androgen receptor (AR) signaling, making this pathway a primary therapeutic target [1]. Consequently, androgen deprivation therapy (ADT), aimed at systemically reducing androgen levels or directly inhibiting AR activity, has long been the cornerstone of treatment for advanced and metastatic disease. While ADT often yields significant initial clinical responses, its efficacy is invariably limited by the eventual emergence of therapeutic resistance. This progression leads to castration-resistant prostate cancer (CRPC), a more aggressive state where the disease advances despite an androgen-depleted environment [1]. The challenge of CRPC spurred the development of next-generation AR-targeted therapies, such as enzalutamide and apalutamide, which provide more potent and comprehensive AR signaling inhibition and became the standard-of-care for this patient population [2, 3]. However, the clinical benefits of these advanced therapies are also frequently curtailed by the development of acquired resistance [4-6]. Addressing this resistance to advanced AR-targeted agents in CRPC is therefore a critical focus of ongoing oncological research and underscores the urgent need for novel therapeutic strategies.

Several molecular mechanisms have been elucidated that confer resistance to AR-targeted therapies. These include, but are not limited to, the restoration of AR-driven transcriptional programs and the circumvention of AR signaling dependency through the activation of alternative transcription factors [7]. Furthermore, emerging evidence has delineated a distinct mechanism known as lineage plasticity, whereby prostate cancer cells escape the luminal epithelial lineage and transform into a multi-lineage, progenitor-like state that is no longer AR-dependent [8-11]. This plasticity has been implicated in regulating therapeutic resistance across diverse human malignancies, including prostate, breast, lung, and pancreatic cancers, as well as melanoma [9, 11-17], and characterized by a spectrum of genomic and transcriptional aberrations [12, 14, 18-32]. For instance, our previous research identified that concurrent loss-of-function alterations in the tumor suppressor genes *TP53* and *RB1* promote treatment resistance in prostate cancer, in part through the activation of Jak-Stat signaling [9, 11]. Intriguingly, many reported mechanisms of therapeutic resistance appear to involve prostate cancer cells acquiring or ‘hijacking’ stem-like, pluripotent characteristics, suggesting that stemness-associated signaling pathways may be directly implicated in processes such as lineage plasticity.

Among these, Notch is a key stemness signaling pathway. The Notch signaling pathway is a highly conserved intercellular communication mechanism crucial for regulating fundamental cellular processes, including cell fate determination, proliferation, differentiation, and apoptosis, which are vital for embryonic development and adult tissue homeostasis [33, 34]. Activation of this pathway occurs through direct cell-to-cell interaction, where in human cells, transmembrane ligands from the Delta-like (DLL1, DLL3, DLL4) and Jagged (JAG1, JAG2) families on one cell engage one of the four Notch receptors (NOTCH1-4) on an adjacent cell. This binding initiates a cascade of proteolytic cleavages, culminating in the release of the Notch intracellular domain. This domain subsequently translocates to the nucleus, where it complexes with the CSL transcription factor and other co-activators to modulate the expression of target genes, such as those in the HES and HEY families (Figure 1). Given its integral role in cellular functions, aberrant Notch signaling is frequently implicated in the pathogenesis of numerous human diseases, including malignancies [35]. Specifically, in prostate adenocarcinoma, the Notch pathway generally exhibits an oncogenic role; activation of Notch signaling has been shown to promote the proliferation, survival, migration, and invasion of prostate cancer cells [36]. On the other hand, Notch signaling exhibits a dramatic functional shift from oncogenic to tumor suppressive during neuroendocrine development from adenocarcinoma of prostate cancer through lineage plasticity [37].

**Figure 1.**
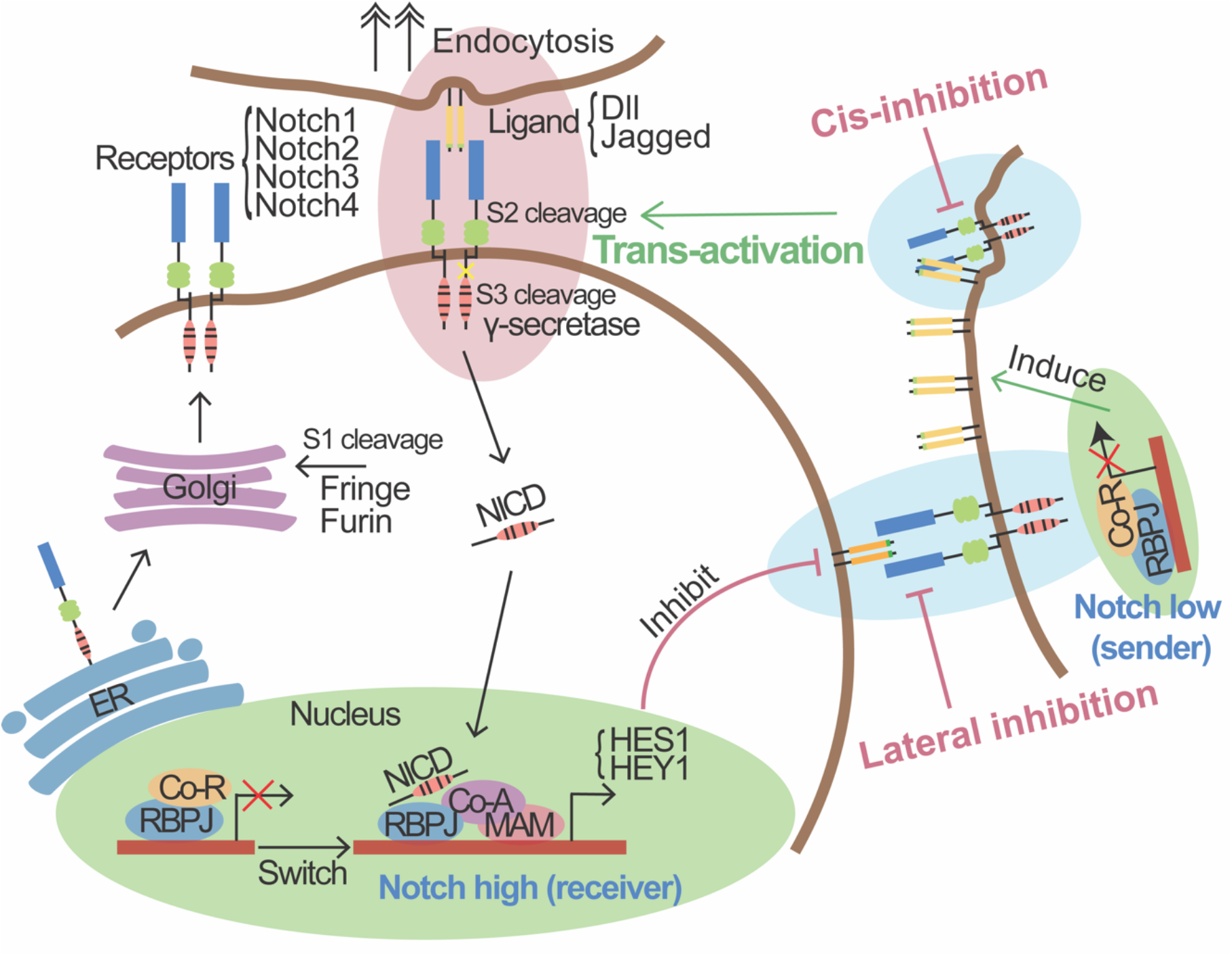
Schematic overview of the Notch signaling pathway. Notch receptors, after S1 cleavage, form heterodimer and traffic to the cell surface. Trans-activation occurs when ligands on an adjacent cell bind a receptor, inducing S2 and S3 cleavages. This releases the NICD, which translocates to the nucleus and converts RBPJ from a transcriptional repressor into an activator by recruiting co-activator (Co-A) proteins like MAM, stimulating target genes such as *HES1* and *HEY1*. This can induce lateral inhibition in neighboring Notch^low^/Ligand^high^ sender cells. Cis-inhibition occurs when ligands and receptors on the same cell interact, attenuating signaling. Co-R: Co-repressor; ER: Endoplasmic Reticulum; MAM: Mastermind-like.

In this study, we demonstrate that activation of Notch signaling is a critical driver of lineage plasticity and is required for resistance to AR-targeted therapy in castration-resistant prostate cancer (CRPC). Furthermore, we show that CRISPR-mediated knockout or targeted inhibition of Notch effectively reverses enzalutamide resistance *in vitro*. Collectively, these findings identify Notch signaling as a crucial executor driving therapy resistance, suggesting it as a promising therapeutic target to overcome resistance to AR-targeted therapies in advanced prostate cancer.

## Results

### Elevated Notch Activity and Expression in Castration-Resistant Prostate Cancer

Notch signaling exhibits inherent heterogeneity across cell populations due to mechanisms such as trans-activation, cis-inhibition, and lateral inhibition (Figure 1). Hence, we started our characterization of Notch signaling at single-cell transcriptomic level including four distinct cell populations: castration-sensitive prostate cancer adenocarcinoma (CSPC^AD^), castration-resistant prostate cancer adenocarcinoma (CRPC^AD^), CRPC progenitor (CRPC^Progenitor^), and CRPC neuroendocrine (CRPC^NE^) (Figure 2A). Visualization of the calculated Notch pathway activity score onto this UMAP revealed a heterogeneous distribution of Notch signaling across these populations, with notably higher activity in majority of the CRPC^AD^ and CRPC^progenitor^ compared to the most naive and early stage treatment-sensitive prostate cancer (CSPC) clusters (Figure 2B). Quantification confirmed the gradual elevation of Notch signaling activity from CSPC to CRPC^AD^ and then to CRPC^Progenitor^. Consistent with previous report, the Notch signaling dropped in the trans-differentiated neuroendocrine prostate cancer cells (Figure 2C). To further determine whether this elevated Notch activity observed during the progression to castration-resistant prostate cancer reflects an intrinsic adaptive response of the luminal epithelial cells-of-origin to androgen deprivation, we then examined Notch signaling dynamics in the wild-type mouse luminal epithelial cells throughout the castration and regeneration cycle [38]. We found that mouse luminal epithelial cells exhibited a progressive increase throughout the castration phase, a period directly analogous to androgen deprivation therapy. Following the reintroduction of androgens, this elevated Notch activity then gradually declined as the prostate underwent regeneration (R2D-R28D) (Figure 2D). To further validate these observations from single-cell analyses with a larger sample number, we analyzed bulk RNA-seq data from human patient samples. This broader dataset corroborated the trend of differential Notch activity observed in the single-cell studies that increase along with cancer progression and resistance but decreased in neuroendocrine prostate cancer cells (Figure 2E).

**Figure 2.**
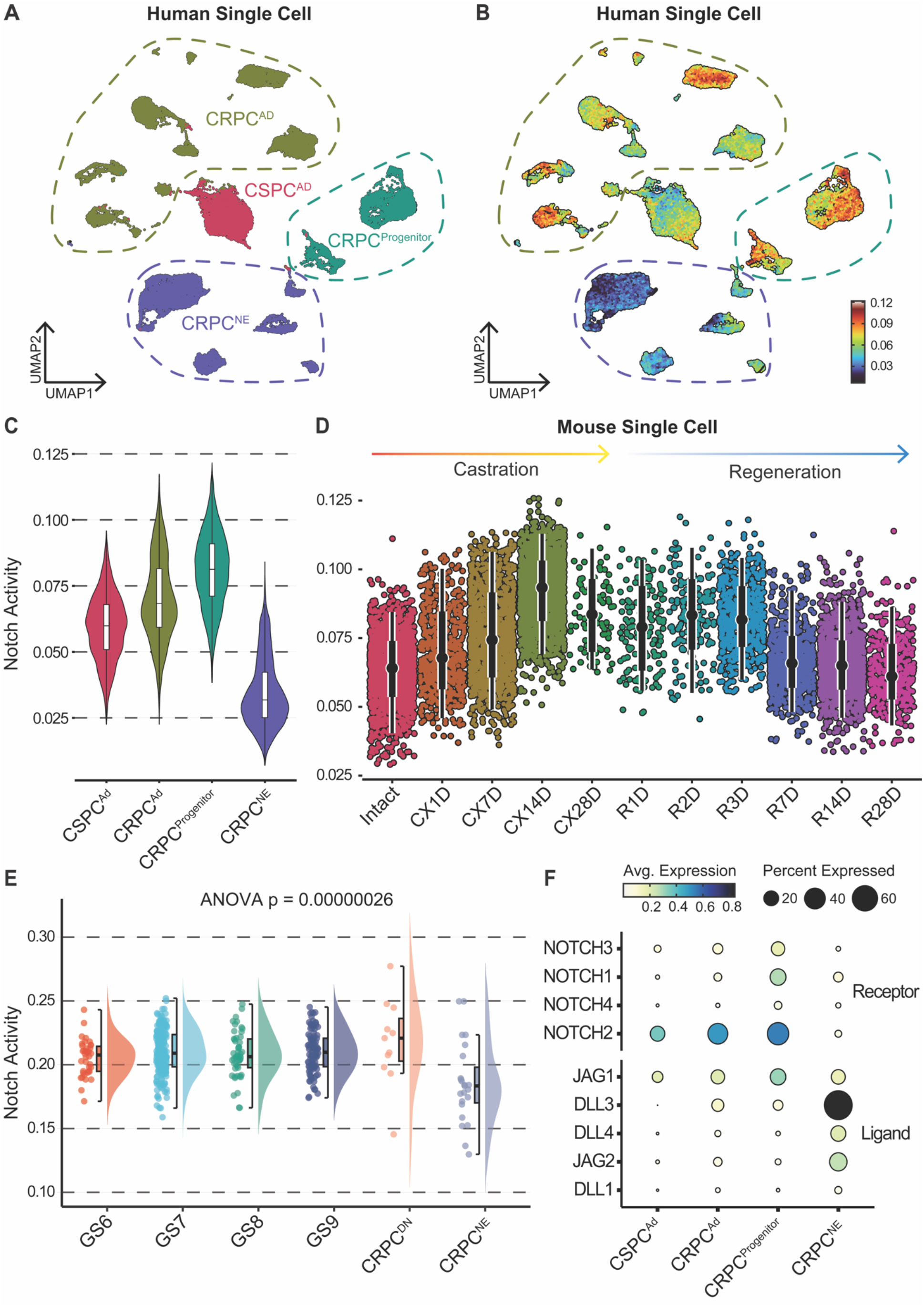
Elevated Notch activity and expression in castration-resistant prostate cancer. (A) UMAP plot of human single-cell RNA sequencing data, illustrating distinct prostate cancer cell populations: castration-sensitive prostate cancer adenocarcinoma (CSPC^AD^), castration-resistant prostate cancer adenocarcinoma (CRPC^AD^), CRPC progenitor (CRPC^Progenitor^), and CRPC neuroendocrine (CRPC^NE^). (B) UMAP plot from (A) colored by Notch activity score, with warmer colors (red/orange) indicating higher activity and cooler colors (blue) indicating lower activity. (C) Violin plots comparing Notch pathway activity scores across the indicated human prostate cancer cell populations. (D) Violin and scatter plots depicting Notch pathway activity in mouse single-cell RNA-seq dataset over a time course of castration (CX1D to CX28D) and subsequent regeneration (R1D to R28D), compared to intact controls. (E) Violin plots showing Notch activity across prostate cancer patient groups. GS: Gleason score; CRPC^DN^: CRPC: AR^-^/NE^-^. (F) Dot plot illustrating the average expression and percentage of cells expressing key Notch receptors and ligands within the distinct human prostate cancer cell populations.

Having established and validated these dynamic changes in overall Notch pathway activity across different disease states and model systems, our next step was to identify the specific Notch signaling components (e.g., ligands, receptors, and target genes) responsible for these alterations by interrogating their expression at the single-cell level within our defined prostate cancer populations. The dot plot (Figure 2F) revealed a distinct and consistent trend in the expression changes of Notch receptors and ligands, aligning with the calculated Notch signaling activity. Three out of four Notch receptors were detected in prostate cancer cells, with their expression increasing progressively from CRPC^AD^ to CRPC^Progenitor^, and then decreasing in CRPC^NE^, as expected. Interestingly, the expression of Notch ligands, including members of the JAG and DLL gene families, exhibited a divergent pattern and were notably upregulated in CRPC^NE^, where Notch receptor expression was lower. This seemingly paradoxical observation is consistent with intrinsic Notch regulatory feedback mechanisms, such as lateral inhibition, wherein active Notch signaling often suppresses ligand expression, confirming the previous characterized Notch activity changes.

### *TP53/RB1* Co-Deficiency Drives Enzalutamide Resistance via Notch Pathway Activation

To explore the mechanisms by which Notch signaling is coupled to prostate cancer progression and lineage plasticity, we first investigated the potential upstream regulators of Notch signaling during this process. Our previous publications indicated that *TP53/RB1* co-deficiency, a common genetic alteration observed in late-stage prostate cancer, can drive treatment resistance and promote lineage plasticity [11]. Based on this, we examined whether the combined loss of *TP53* and *RB1* could also activate Notch signaling. To address this, we assessed the expression of Notch pathway genes following *TP53/RB1* knockdown in prostate cancer cells. As shown in Figure 3A, induction of *TP53/RB1* knockdown (gTP53/RB1 + Dox) significantly increased the expression of *NOTCH1* and its downstream target gene *HES1*.

**Figure 3.**
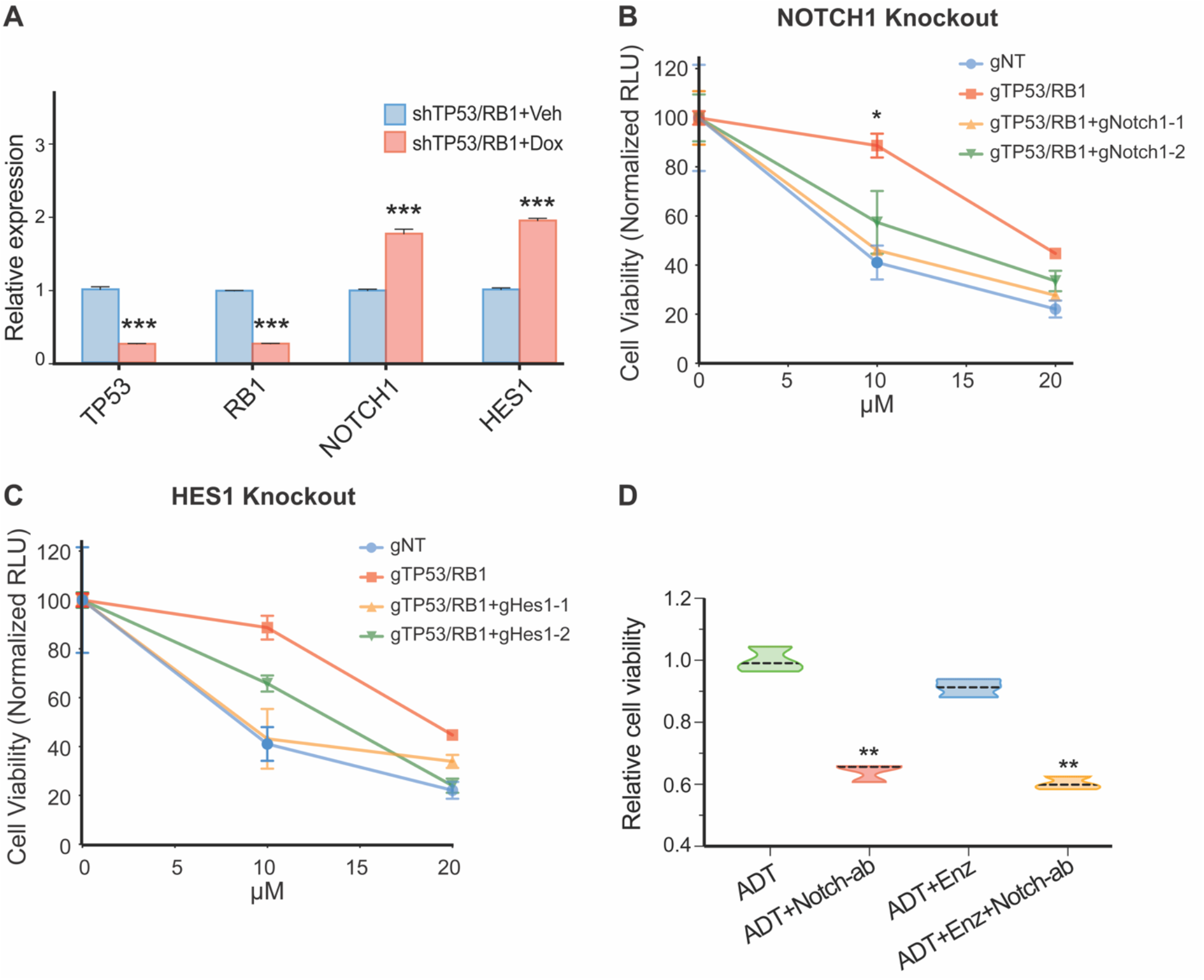
Notch1 as the top candidate effector of *TP53/RB1* co-deficiency and enzalutamide resistance. (A) Relative gene expression level of *TP53/RB1* downstream effectors in cells transduced with an inducible KD system. p-values were calculated by multiple t-test. For all panels, mean ± s.e.m. is represented and *** represents p<0.001. (B, C) Relative cell viability as measured by Cell Titer-Glo assay, showing cells transduced by guide RNAs targeting annotated genes. p-values were calculated by one-way ANOVA. For all panels, mean ± s.e.m. is represented and * represents p<0.05. (D) Relative cell viability as measured by Cell Titer-Glo assay, showing gTP53/RB1-ER cells treated with different combination of drugs. ADT represents androgen deprivation therapy which is achieved by CSS medium. Notch-ab represents Notch antibody. p-value were calculated by one-way ANOVA. For all panels, mean ± s.e.m. is represented and ** represents p<0.01.

Having established the upregulation of *NOTCH1* and its target *HES1* downstream of *TP53*^*-*^ */RB1*^*-*^, we next investigated whether Notch signaling is a driver of treatment resistance in TP53/RB1-deficient prostate cancer cells. We utilized CRISPR-Cas9 to individually knock-out *NOTCH1* or *HES1* genes and followed by cell viability assay in the condition of Enzalutamide treatment. Consistent with our previous report, control cells transduced with a non-targeting guide RNA (gNT) exhibited sensitivity to enzalutamide, whereas *TP53/RB1* double knockout (gTP53/RB1) cells showed significantly higher viability across increasing concentrations of the drug (Figures 3B and 3C). Notably, knockout of *NOTCH1* using two independent guide RNAs (gNotch1-1 and gNotch1-2) in the gTP53/RB1 background almost completely reversed resistance, re-sensitizing these cancer cells to enzalutamide treatment to a level comparable with gNT controls (Figure 3B). A similar re-sensitization to enzalutamide was observed upon knockout of the *NOTCH1* downstream effector *HES1* using two distinct guide RNAs (gHes1-1 and gHes1-2) (Figure 3C). These data indicate that, beyond mere association, Notch signaling, including its downstream effector *HES1*, is functionally involved in prostate cancer resistance as a crucial driver mechanism.

### Notch Signaling is a Druggable Target

Although concurrent alterations in the *TP53* and *RB1* loci have been implicated in conferring lineage plasticity and therapy resistance, direct pharmacologic activation of *TP53* and *RB1* is not currently feasible [11]. This led us to investigate an alternative approach: inhibiting Notch signaling to re-sensitize prostate cancer to enzalutamide treatment in our preclinical model. While γ-secretase inhibitors (GSIs), small molecules that broadly inhibit Notch signaling, have shown preclinical promise, their clinical development faces challenges due to mechanism-based toxicities arising from non-selective γ-secretase inhibition [39]. We therefore hypothesized that a Notch1-specific antibody (Notch1-ab) could provide a more targeted and potentially safer therapeutic strategy to inhibit Notch signaling.

To create a robust model for this, we first aimed to develop a stably enzalutamide-resistant cell line from the gTP53/RB1 LNCaP/AR cells, which initially exist in a multi-lineage, plastic state with the potential to move back to AR dependency if the selection pressure is removed. These gTP53/RB1 cells were subjected to prolonged treatment with a low dosage of enzalutamide for six months to maintain selection pressure. This long-term culture under enzalutamide treatment resulted in the establishment of a derivative cell line, designated gTP53/RB1-ER (Enzalutamide Resistant). These gTP53/RB1-ER cells exhibited stable and continuous upregulation of canonical neuroendocrine lineage markers, confirming the development of a stable, therapy-resistant phenotype suitable for testing efficacy of Notch inhibitor.

We then assessed the impact of a NOTCH1 inhibitor, LEAF™ purified anti-human Notch1 antibody (Notch-ab), on the viability of these gTP53/RB1-ER cells, alone or in combination with enzalutamide, under androgen deprivation therapy (ADT) conditions (CSS medium). As measured by cell viability assay through CellTiter-Glo, the gTP53/RB1-ER cells were highly resistant to enzalutamide based on similar cell viability between ADT+Enz and ADT alone (Figure 3D). However, treatment with the Notch1 antibody significantly reduced cell viability (ADT+Notch-ab vs. ADT, p<0.01). Notably, when the Notch1 antibody was combined with enzalutamide (ADT+Enz+Notch-ab), cell viability was reduced to a similar extent as with the Notch antibody alone, and significantly more than enzalutamide treatment alone (Figure 3D). These findings demonstrate that pharmacological inhibition of Notch can overcome established enzalutamide resistance in this stably resistant, lineage-altered prostate cancer cell model, providing a strong rationale for further *in vivo* investigation.

## Discussion

The development of resistance to AR-targeted therapies represents one of the most formidable challenges in advanced prostate cancer management. While the clinical efficacy of next-generation AR antagonists like enzalutamide has transformed treatment paradigms for castration-resistant prostate cancer (CRPC), the inevitable emergence of therapeutic resistance continues to drive disease progression and limit patient survival. Our findings reveal critical mechanistic insights into this resistance phenomenon by identifying Notch signaling as a key downstream effector of *TP53/RB1* co-deficiency that drives lineage plasticity and confers enzalutamide resistance in CRPC. Specifically, we demonstrate that Notch activity is elevated across prostate cancer progression and acquiring resistance with the highest expression in therapy-resistant AR-/NE-progenitor cell states and followed by striking decrease in transdifferentiated neuroendocrine prostate cancer stage. Functionally, CRISPR-mediated knockout of either *NOTCH1* or *HES1* effectively reversed enzalutamide resistance in TP53/RB1-deficient cells. Furthermore, pharmacological inhibition of Notch using a blocking antibody overcome established resistance in stably enzalutamide-resistant cell lines, demonstrating the therapeutic potential of targeting this pathway.

Although the Notch pathway is a canonical regulator of stemness and has been previously characterized across various prostate cancer subtypes, our study offers a comprehensive, full-spectrum analysis of dynamic changes in Notch signaling activity. Notably, we newly identified a peak in Notch activity within the double-negative CRPC subtype enriched for progenitor-like SOX2+ cells, aligning with Notch’s established role in stem cell and development. Furthermore, our findings address critical gaps in the clinical targeting of Notch. While earlier studies associated NOTCH overexpression with aggressive disease features and poor prognosis in primary prostate adenocarcinoma, which suggest potential benefit from Notch inhibition, our results provide direct preclinical evidence supporting this strategy. Specifically, dual inhibition of NOTCH and its downstream effector HES reverses enzalutamide resistance through both genetic knockout and antibody-based approaches.

Collectively, our study establishes Notch signaling as a crucial and targetable executor of lineage plasticity and enzalutamide resistance in TP53/RB1-deficient prostate cancer (Figure 4). These *in vitro* findings provide a strong rationale for subsequent translational work. Future efforts will focus on validating the efficacy of NOTCH inhibitors to overcome lineage plasticity-driven resistance *in vivo* using relevant preclinical models. Furthermore, we plan to extend these investigations to diverse prostate cancer models beyond LNCaP/AR derivatives to ascertain the broader applicability of targeting this pathway. Such studies, alongside efforts to identify predictive biomarkers for Notch dependency, will be essential for advancing this approach towards clinical application for patients battling therapy-resistant prostate cancer.

**Figure 4.**
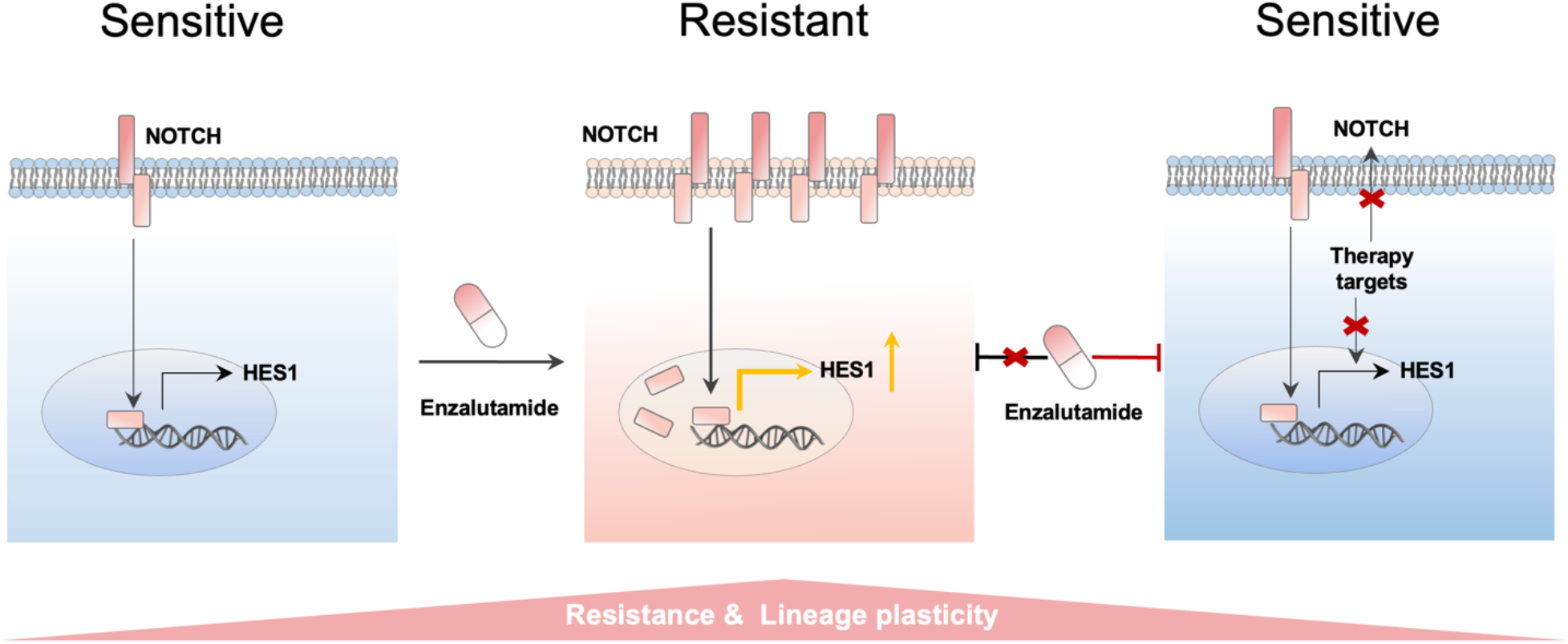
Model for Notch-mediated enzalutamide resistance and re-sensitization. (Left) Treatment-sensitive cells exhibit baseline Notch receptor activity and *HES1* expression. (Middle) Upon Enzalutamide treatment, resistant cells show upregulated Notch receptor expression and elevated *HES1*, associated with resistance and lineage plasticity. (Right) Co-administration of a Notch inhibitor with Enzalutamide blocks aberrant Notch-HES1 signaling, restoring treatment sensitivity.

## Methods

### Cell Lines and Culture Conditions

The LNCaP/AR prostate cancer cell line (obtained from the laboratory of C. Sawyers, MSKCC) was used as the parental line for generating experimental models of this study. LNCaP/AR cells were cultured in RPMI 1640 medium supplemented with 10% fetal bovine serum (FBS), 1% L-glutamine, 1% penicillin–streptomycin, 1% HEPES, and 1% sodium pyruvate. Cell cultures were maintained at 37°C in a humidified atmosphere containing 5% CO2 and routinely assessed for mycoplasma contamination using the MycoAlert Plus Mycoplasma Detection kit (Lonza, LT07-710), with all results being negative. Cell line identity was validated annually through human short tandem repeat profiling.

The *TP53* and *RB1* co-deficient (gTP53/RB1) LNCaP/AR cell line was generated using CRISPR-Cas9 technology as described below. For experiments involving enzalutamide treatment, gTP53/RB1 cells were cultured in RPMI 1640 medium supplemented with 10% charcoal-stripped serum (CSS medium; to maintain androgen-deprived conditions). For *TP53/RB1* knockdown induction, cells were treated with doxycycline (Dox) at 1 µg/mL for 72 hours prior to analysis. Vehicle-treated cells received an equivalent volume of Dimethyl sulfoxide (DMSO). The stably resistant gTP53/RB1-ER cell line is generated by culturing gTP53/RB1 cells in CSS medium and treating with a low dosage of enzalutamide (5 µM) for a period of 6 months.

### CRISPR and shRNA

Lentiviral-based constructs were used for CRISPR-Cas9-mediated knockout of *TP53, RB1* (for generating the initial gTP53/RB1 line), *NOTCH1*, and *HES1* in LNCaP/AR cells. LNCaP/AR cells were seeded at 400,000 cells per well in 6-well plates. The next day, medium was replaced with medium containing 50% virus, 50% fresh culture medium, and 5 μg/mL polybrene. After 24 hours, the virus-containing medium was replaced with normal culture medium. Cells were selected with 2 μg/mL puromycin for 4 days or 5 μg/mL blasticidin for 5 days. The All-In-One lentiCRISPRv2 (Addgene plasmid 52961), LentiCRISPRv2GFP (Addgene plasmid 82416), LentiCRISPRv2-mCherry (Addgene plasmid 99154), pLKO5.sgRNA.EFS.RFP (Addgene plasmid 57823), pLKO5.sgRNA.EFS.GFP, and lentiCas9-Blast (Addgene plasmid 52962) plasmids were used to generate the CRISPR and guide RNAs. Guide RNAs were designed using the Benchling guide RNA designing tool. Non-targeting guide RNA (sgNT) constructs with an empty space holder served as controls. shRNA construct LT3GEPIR (pRRL-TRE3G-GFP-miRE-PGK-PuroR-IRES-rtTA3) was originally obtained from the laboratory of J. Zuber at the Research Institute of Molecular Pathology.

### RNA Extraction and Quantitative Real-Time PCR (qPCR)

Total RNA was extracted using Trizol from LNCaP/AR cells with or without doxycycline induced *TP53* and *RB1* knockdown. cDNA was synthesized from 200 ng of RNA template using the SuperScript IV VILO Master Mix with ezDNase enzyme (Thermo Fisher, 11766500). qPCR was performed using 2x PowerUp SYBR Green Master Mix (Thermo Fisher, A25778) on a QuantStudio Real-Time PCR (qPCR) system. Primers for human *NOTCH1, HES1, TP53, RB1*, and housekeeping gene *GAPDH* were used.

### Cell Viability Assay and Enzalutamide Treatment

Cell viability was assessed using the CellTiter-Glo® Luminescent Cell Viability Assay (Promega, G7570) according to the manufacturer’s protocol, with luminescence quantified on a SpectraMax iD3 automatic plate reader. All viability data were normalized to the respective vehicle-treated control cells for each cell line or condition.

*Enzalutamide Dose-Response Curves:* LNCaP/AR-derived cells, including non-targeting control (sgNT), *TP53/RB1* co-deficient (gTP53/RB1), gTP53/RB1 with *NOTCH1* knockout (gTP53/RB1+sgNotch1-1 and gTP53/RB1+sgNotch1-2), and gTP53/RB1 with *HES1* knockout (gTP53/RB1+sgHes1-1 and gTP53/RB1+sgHes1-2), were seeded at 4,000 cells per well in 96-well plates in CSS medium. After 24 hours, cells were treated with 0, 1, 5, 10, and 20 µM of enzalutamide (sourced from the Organic Synthesis Core Facility at MSKCC) or vehicle control (DMSO) for 8 days.

*Notch Inhibitor Assays:* The stably enzalutamide-resistant gTP53/RB1-ER cells were seeded at 4,000 cells per well in 96-well plates in CSS medium. After 24 hours, cells were treated with 10 µM enzalutamide, 10 µM LEAF™ purified anti-human Notch1 antibody (Biolegend, #352111), or a combination of both, as indicated in Figure 3D. Control groups received DMSO as vehicle. Treatments were maintained for 8 days.

### Bioinformatics

Data analysis was performed using R. Gene expression data (counts and Transcripts Per Million -TPMs) were initially loaded. The MSigDB Hallmark Notch Signaling gene set was utilized as a defined gene signature. Human gene symbols were converted to mouse orthologs where necessary using nichenetr (https://github.com/saezlab/nichenetr). Single-cell RNA sequencing (scRNA-seq) data from the Human Prostate Single-cell Atlas (HuPSA), Mouse Prostate Single-cell Atlas (MoPSA), and C42B xenograft datasets were analyzed using the Seurat V5 (https://github.com/satijalab/seurat). Standard scRNA-seq workflows included data normalization (LogNormalize), scaling, principal component analysis (PCA), and Uniform Manifold Approximation and Projection (UMAP) for dimensionality reduction. Batch correction and data integration across different studies or samples were carried out using Harmony (https://github.com/immunogenomics/harmony) as implemented within Seurat’s IntegrateLayers function. Notch pathway activity and other gene signature scores were calculated per cell using UCell (https://github.com/carmonalab/UCell). For bulk RNA-seq datasets like ProAtlas, pathway activity scores were derived using singscore (https://bioconductor.org/packages/singscore/) from TPM values. Data manipulation and wrangling were facilitated by dplyr (https://github.com/tidyverse/dplyr) and tidyr (https://github.com/tidyverse/tidyr). Visualization of results, including PCA plots, boxplots, violin plots, feature plots, dot plots, and heatmaps, was performed using ggplot2 (https://github.com/tidyverse/ggplot2), pheatmap (https://github.com/raivokolde/pheatmap), gghalves (https://github.com/erocoar/gghalves), and SCpubr (https://github.com/enblacar/SCpubr) for specialized single-cell visualizations.

## Statistical Analysis

Data are presented as mean ± s.e.m. from at least three independent experiments. Statistical significance for qPCR data was assessed using multiple t-tests. For cell viability assays, one-way ANOVA followed by Bonferroni or Benjamini correction were performed to determine significance at each drug concentration or each treatment condition. P-values for dose-response curves were calculated by non-linear regression with an extra sum-of-squares F-test. A p-value < 0.05 was considered statistically significant (*p<0.05, **p<0.01, ***p<0.001, ****p<0.0001). Statistical analyses and graphing were performed using GraphPad Prism (V9.3.1).

## Acknowledgments

This work was supported or partially supported by: National Cancer Institute/National Institutes of Health: R00CA218885, R37CA258730, R01CA288820, R01CA292949 P. Mu; Department of Defense: W81XWH-18-1-0411 and W81XWH21-1-0520 P. Mu; Cancer Prevention Research Institute (CPRIT): RR170050, RP220473, P. Mu, Prostate Cancer Foundation: 17YOUN12 P. Mu, Yale Cancer Center CCSG Pilot Grant. P30CA016359 P. Mu. and I.Y.K.

## CRediT author statement

**Yuyin Jiang**: Conceptualization, Visualization, Writing-Original draft preparation; **Siyuan Cheng**: Conceptualization, Methodology, Software Data curation; **Longjun Li**: Visualization; **Melanie Fraidenburg**: Validation**; Isaac Kim**: Supervision; **Su Deng**: Methodology, Validation, Supervision; ***Ping Mu***: Supervision, Writing-Reviewing and Editing.

## Data and code availability

The R codes necessary for reproducing all bioinformatics figures are available at https://github.com/schoo7/notch. The single-cell RNA sequencing data HuPSA and MoPSA data are downloaded from figShare: https://doi.org/10.1038/s41698-024-00667-x [40]. The bulk RNA sequencing data from patient (ProAtlas) generated by our previous publication is available through https://pcatools.shinyapps.io/HuPSA-MoPSA/ [40].

